# SEQUENCE VARIATIONS ASSOCIATED WITH NOVEL PURPLE-PERICARP SUPER-SWEETCORN COMPARED TO ITS PURPLE-PERICARP MAIZE AND WHITE SUPER-SWEETCORN PARENTS

**DOI:** 10.1101/2022.07.28.501808

**Authors:** Apurba Anirban, Ardashir Kharabian Masouleh, Robert J Henry, Tim J O’Hare

## Abstract

Recently, a novel purple-pericarp super-sweetcorn line, ‘Tim1’ (*A1A1*.*sh2sh2*) is derived from the purple-pericarp maize ‘Costa Rica’ (*A1Sh2*.*A1Sh2*) and white *shrunken2* (*sh2*) super-sweetcorn ‘Tims-white’ (*a1sh2*.*a1sh2*), however information regarding purple colour controlling anthocyanin biosynthesis genes and sweetcorn gene is lacking. Specific sequence differences in the CDS (coding DNA sequence) and promoter regions of the anthocyanin biosynthesis structural genes, *anthocyanin1* (*A1*), *purple aleurone1* (*Pr1*) and regulatory genes, *purple plant1* (*Pl1*), *plant colour1* (*B1*), *coloured1* (*R1*), and the sweetcorn structural gene, *shrunken2* (*sh2*) were investigated using the publicly available annotated yellow starchy maize, B73 (NAM5.0) as a reference genome. In the CDS region, the *A1, Pl1* and *R1* gene sequence differences of ‘Tim1’ and ‘Costa Rica’ were similar, as they control purple-pericarp pigmentation, however the *B1* gene showed similarity between the ‘Tim1’ and ‘Tims-white’ lines, which may indicate that it does not have a role in controlling pericarp colour, unlike a previous study. In case of *Pr1* gene, unlike ‘Costa Rica’, 6- and 8-bp dinucleotide (TA) repeats were observed in the promoter region of the ‘Tims-white’ and ‘Tim1’ lines, respectively, indicating the defective functionality (redder colour in ‘Tim1’ than purple in ‘Costa Rica’) of the recessive *pr1* allele. In the sweetcorn structural gene (*sh2*), sequence similarity was observed between purple-sweet ‘Tim1’ and its white-sweet parent ‘Tims-white’, as both display a shrunken phenotype in their mature kernels. These findings revealed that the developed purple-sweet line is different than the reference yellow-nonsweet line regarding both the anthocyanin biosynthesis and sweetcorn genes.

## Introduction

There is no previous scientific report available on the development of purple-pericarp super-sweetcorn based on the *shrunken2* super-sweet gene because of the tight genetic linkage between an anthocyanin biosynthesis inactive allele, *anthocyaninless1 (a1)*, and super-sweet allele *shrunken2* (*sh2)* (Civardi et al., 1994, Yao et al., 2002). Recently, a novel purple-pericarp super-sweetcorn F3 lines was developed from the cross between purple-pericarp maize parent, ‘Costa Rica’ and white super-sweetcorn parent, ‘Tims-white’ followed by breaking the close genetic linkage (Anirban and O’Hare, 2020). Later, a F6 fixed line, ‘Tim1’ was developed. Field experiments revealed that the purple-pericarp maize ‘Costa Rica’ and the purple-pericarp super-sweetcorn ‘Tim1’ were both homozygous regarding the dominant allele of the anthocyanin biosynthesis gene, *A1* (*anthocyanin1*). In the same way, the white super-sweetcorn line, ‘Tims-white’ and the novel purple-pericarp super-sweetcorn line, ‘Tim1’ were both homozygous for the recessive allele of the sweetcorn mutation allele, *sh2*. The presence of *A1* and *sh2* has been confirmed by only visual phenotyping. The genetic linkage between them is confirmed by both field observation and genomic analysis (Anirban et al., 2022). However, no genomic information of this novel genotype is available regarding the anthocyanin biosynthesis structural and regulatory genes, along with sweetcorn structural gene.

Regulatory genes known as transcription factors are also required for anthocyanin pigmentation (Petroni et al., 2014; Petroni and Tonelli, 2011) along with the anthocyanin biosynthesis structural gene, for example, *A1*. The Myb transcription factor *Pl1 (purple plant1)* and *C1 (coloured aleurone1* control pigmentation in the pericarp and aleurone layers of the cob kernel, respectively (Procissi et al., 1997). On the other hand, the bHLH transcription factor *B1* (*plant colour1*) is responsible for plant colour development, while the *R1 (coloured1)* gene play a vital role in pericarp and aleurone pigmentation (Chatham and Juvik, 2021; Chatham et al., 2019). To develop cyanidin (purple pigment) or pelargonidin (red pigment) based anthocyanin, expression of anthocyanin biosynthesis another structural gene, *Pr1 (purple/red aleurone1)* is also required.

Study by Schwarz-Sommer et al., (1987) found one frame-shift mutation in the CDS region of the *A1* gene, which inactivated its functionality. In comparison, the *B1* gene has been reported to have allelic sequence diversity in the promoter region (Radicella et al., 1992). Previous study by Cone et al., (1993) also reported no difference between CDS region of dominant and recessive alleles of the *Pl1* gene, however sequence difference was observed in the promoter region. Similarly, different *Pr1* alleles have shown mutations in the promoter region (Sharma et al., 2011). In contrast, alleles of the *R1* gene have been reported to have differences in the coding region (Consonni et al., 1993). Recently, sweetness induced by the recessive allele of the *sh2* gene has been attributed to a single SNP in the coding region (Ruanjaichon et al., 2021). In summary, the above studies reported that *A1, R1* and *sh2* genes have mutations in the coding region, while *B1, Pl1* and *Pr1* genes have mutations in the promotor regions.

The objectives of this investigation were to identify SNP (single-nucleotide polymorphism) and InDel (insertion/ deletion mutations) in the purple-pericarp super-sweetcorn line (‘Tim1’) in comparison to its purple-pericarp maize parent (‘Costa Rica’) and white super-sweetcorn parent (‘Tims-white’) within the CDS and promoter regions of structural anthocyanin biosynthesis genes, *A1* and *Pr1*; the regulatory Myb transcription factor gene, *Pl1*; the bHLH transcription factor genes, *B1* and *R1*; and the sweetcorn structural gene, *sh2*, to identify allele similarities in common with each parent, that may potentially be functional.

## Results

### Identification of SNPs and InDels in the CDS region of target genes

Non-synonymous SNPs leading to amino acid change in the coding DNA sequence (CDS) region of anthocyanin biosynthesis structural gene *A1 (anthocyanin1)* and *Pr1* (*purple/red aleurone1*), regulatory anthocyanin biosynthesis Myb transcription factor gene *Pl1 (purple plant1)* and bHLH transcription factor genes *R1 (coloured1)* and *B1 (plant colour1)* and the sweetcorn structural gene, *sh2 (shrunken2)* were investigated (Table 1).

**Table 1.**
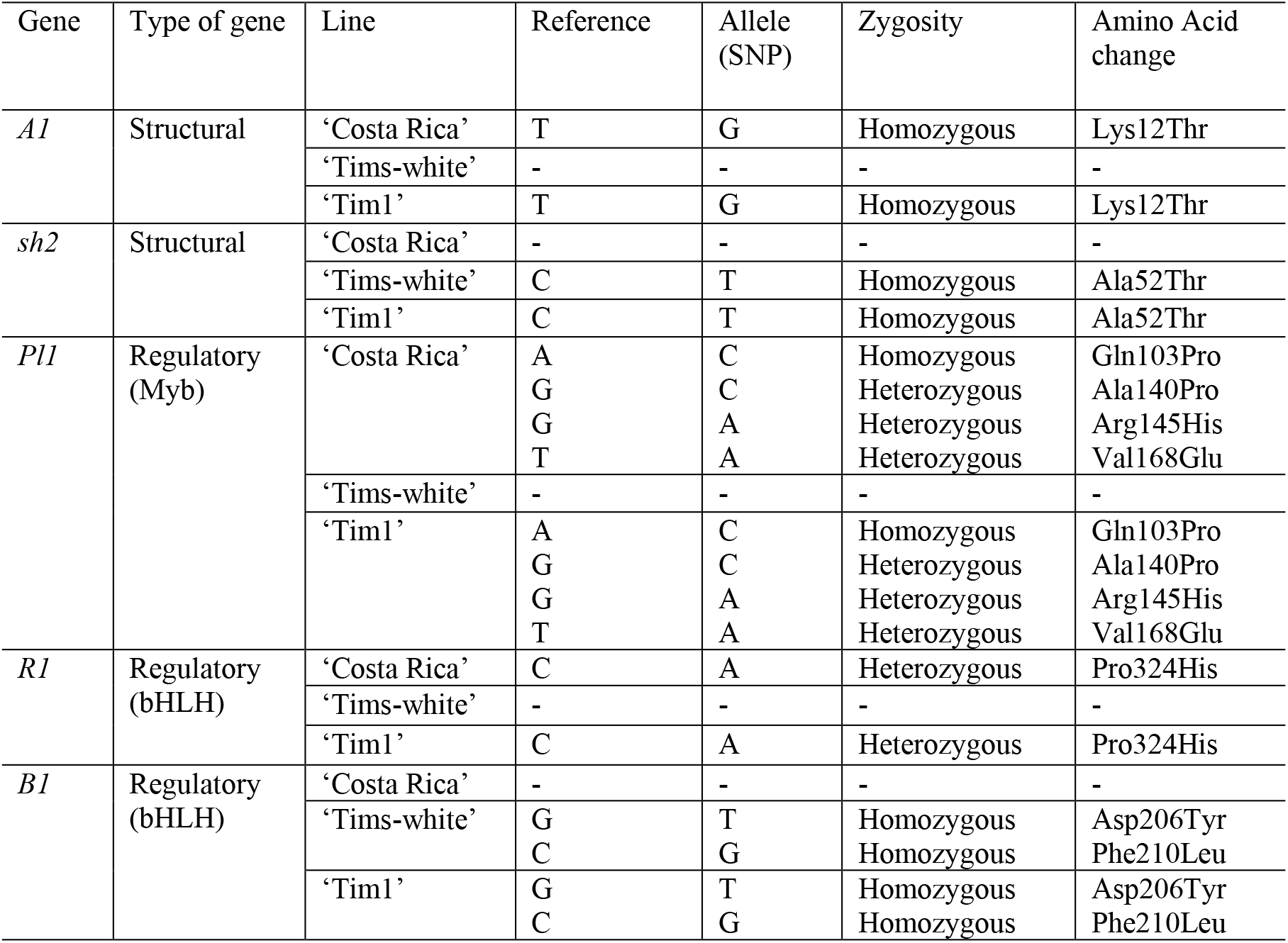
The nonsynonymous SNPs in the CDS region of target genes of ‘Tim1’ in common with ‘Costa Rica’ or ‘Tims-white’ parent. Dash (-) means there is no SNP present.

A SNP that changes a codon resulting in the same amino acid is referred to as a synonymous SNP, while a SNP that changes a codon that produces a different amino acid is referred to as a nonsynonymous SNP, and are considered as a functional SNP (Kharabian, 2010). SNP having same nucleotide in both homologous chromosome and different from the reference genome is known as homozygous SNP, and SNP having different nucleotide in the homologous chromosome and one match with the reference genome and other differ is considered as heterozygous SNP (Wang et al., 2020).

### Anthocyanin biosynthesis structural gene, *A1 (anthocyanin1)*

A single homozygous nonsynonymous SNP (T→G) (Table 1, Fig 1A) in the CDS region of the anthocyanin biosynthesis structural gene, *A1*, of the ‘Costa Rica’ parent, and developed line, ‘Tim1’ was observed, which is responsible for the change of amino acid into threonine than lysine in the annotated B73 reference. By contrast, no nonsynonymous SNPs in the ‘Tims-white’ parental line were detected.

**Fig 1.**
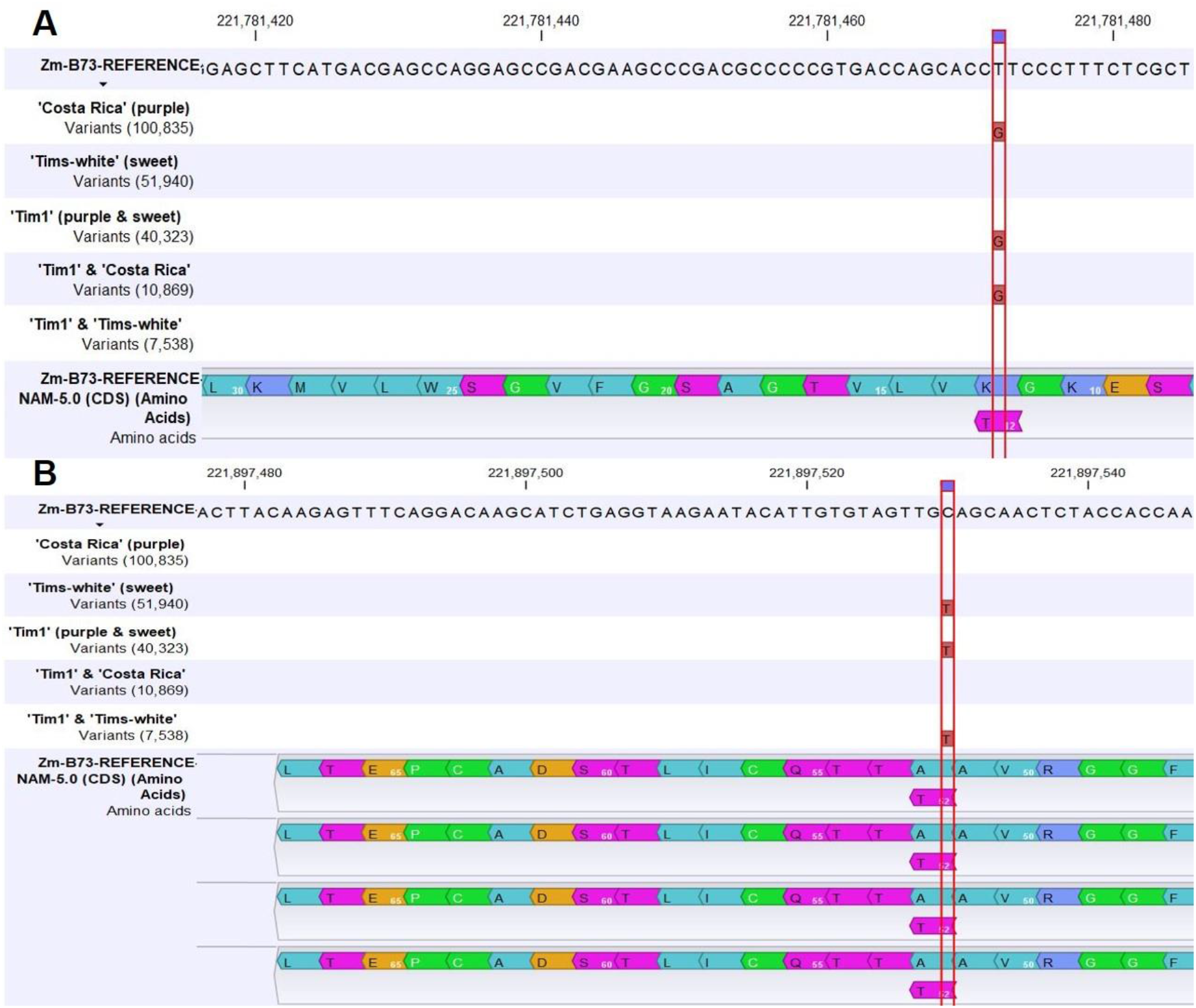
SNPs in the structural genes: **A)** A single homozygous non-synonymous SNP in ‘Tim1’ and ‘Costa Rica’ in the structural gene *A1* (*anthocyanin1*), and **B)** A single homozygous non-synonymous SNP between ‘Tim1’ and ‘Tims-white’ in the structural gene *sh2* (*shrunken2*).

### Anthocyanin biosynthesis structural gene, *Pr1* (*purple/red aleurone1*)

In the case of another anthocyanin biosynthesis structural gene *Pr1* (*purple/red aleurone1*), which determines whether the purple colour will be cyanidin (deep purple) or pelargonidin (reddish purple) there was no nonsynonymous SNP found common between ‘Tim1’ and ‘Costa Rica’ or ‘Tim1’ and ‘Tims-white’ in the CDS region.

### Sweetcorn structural gene, *sh2* (*shrunken2*)

For the supersweet mutation, *sh2*, a single homozygous nonsynonymous SNP (C→T) (Fig 1B) in the CDS region of the purple-pericarp super-sweetcorn line, ‘Tim1’ and the white super-sweetcorn parent ‘Tims-white’ was observed, which is responsible for the change of amino acid into threonine compared to alanine in the annotated B73 reference. There were no nonsynonymous SNPs observed in the non-sweet purple maize parent, ‘Costa Rica’.

### Anthocyanin biosynthesis regulatory gene, *Pl1* (*purple plant1*)

Regarding the regulatory *Pl1* gene, three common nonsynonymous SNPs at CDS 1 and CDS 2 of the reference genome and an additional non-synonymous SNPs at CDS 2 was observed in ‘Costa Rica’ parent and ‘Tim1’, and no nonsynonymous SNPs at these positions within the CDS in ‘Tims-white’ parent were observed.

Out of four SNPs, one homozygous (Fig 2A) nonsynonymous SNP (A→C) in ‘Costa Rica’ and ‘Tim1’ was detected, that changed the amino acid into proline, when compared to the reference amino acid, glutamine (Table 1). There were also three heterozygous (Fig 2B) nonsynonymous SNPs of the *Pl1* gene present in ‘Costa Rica’ parent. The same three heterozygous non-synonymous SNPs at the same genomic position were also present in the developed ‘Tim1’ line. In addition, a six base pair deletion (Fig 2C) in the CDS region was also observed in both ‘Costa Rica’ and ‘Tim1’.

**Fig 2.**
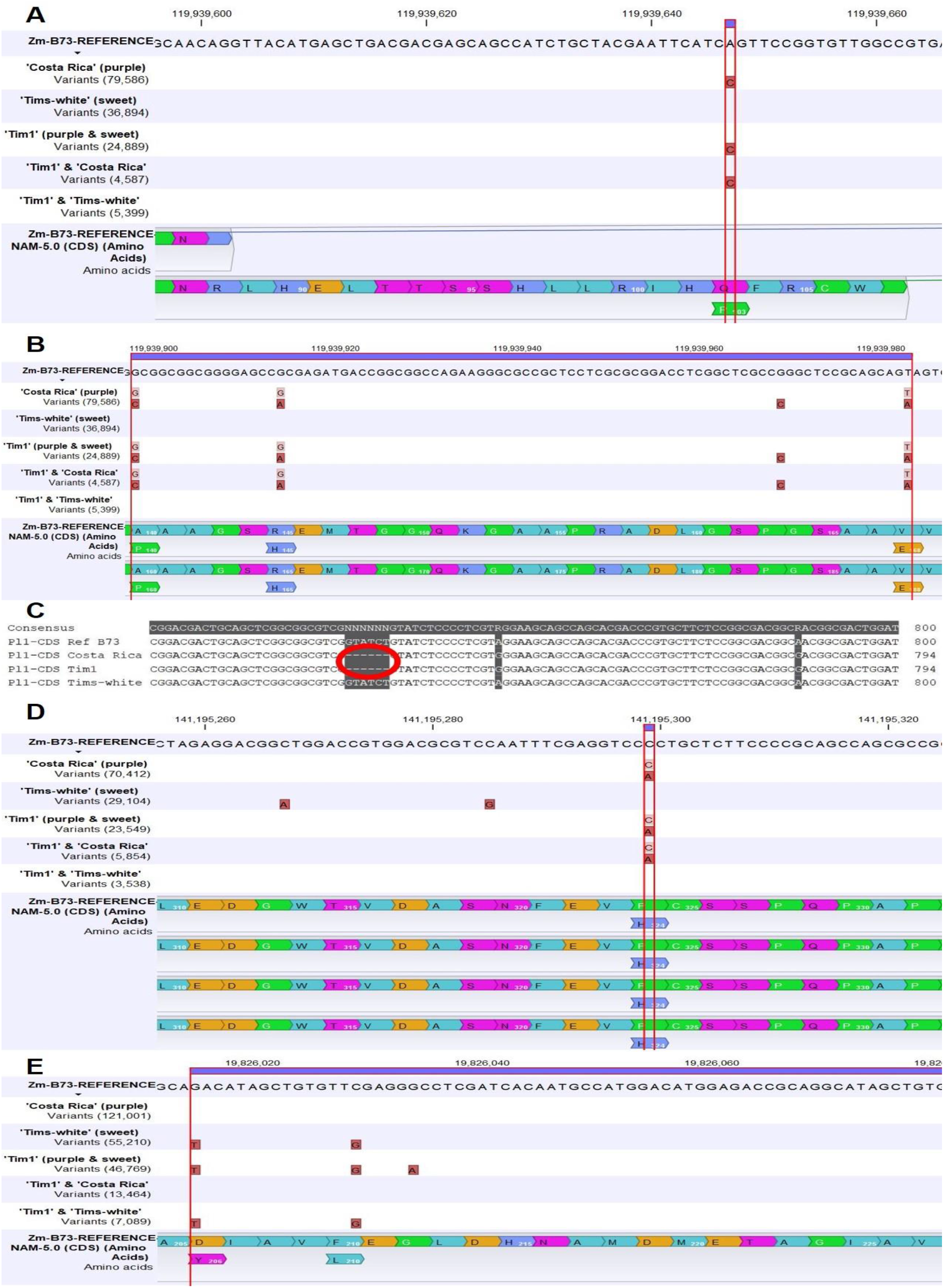
SNPs in the regulatory genes: **A)** A single homozygous nonsynonymous SNP common between ‘Tim1’ and ‘Costa Rica’ in the regulatory gene *Pl1* (*purple plant1*); **B)** Three common heterozygous nonsynonymous SNPs between ‘Tim1’ and ‘Costa Rica’ in the regulatory gene *Pl1*; **C)** A six bp deletion in the CDS region of both in ‘Tim1’ and ‘Costa Rica’ in the regulatory gene *Pl1*, Deletion is indicated by a red circle; **D)** A single heterozygous nonsynonymous SNP common between ‘Tim1’ and ‘Costa Rica’ in the regulatory gene *R1* (*coloured1*); and **E)** Two common homozygous nonsynonymous SNPs between ‘Tim1’ and ‘Tims-white’ in the regulatory gene *B1* (*plant colour1*).

### Anthocyanin biosynthesis regulatory gene, *R1* (*coloured1*)

In the case of the *R1* gene, one heterozygous nonsynonymous SNP (C→A) was found in both ‘Costa Rica’ parent and ‘Tim1’ progeny, and it changed the amino acid into histidine than present in the reference as proline (Table 1, Fig 2D). There was no nonsynonymous SNP observed in the ‘Tims-white’ parental line.

### Anthocyanin biosynthesis regulatory gene, *B1* (*plant colour1*)

Regarding the *B1* gene, two common homozygous nonsynonymous SNPs in the ‘Tims-white’ parent and the developed ‘Tim1’ lines were present, and no nonsynonymous SNP was found in ‘Costa Rica’ parent at those positions in the CDS region (Table 1, Fig 2E).

### Identification of SNPs and InDels in the promoter region of target genes

SNPs and InDels in the promoter region of the target genes were investigated to identify similarity of ‘Tim1’ with each parent that may potentially determine the phenotypes observed.

In the promoter region of the *A1* gene (up to 1.5 kb 5’ upstream region), there were 14 SNPs common to ‘Costa Rica’ parent and ‘Tim1’ progeny (Table 2) when compared to the B73 reference genome. There were also two insertions, one of 1 bp and one of 2 bp. Four deletions were also observed in the promoter region (2 bp, 3 bp, 1 bp and 11 bp). By contrast the ‘Tims-white’ parental line was identical to the reference genome.

**Table 2.**
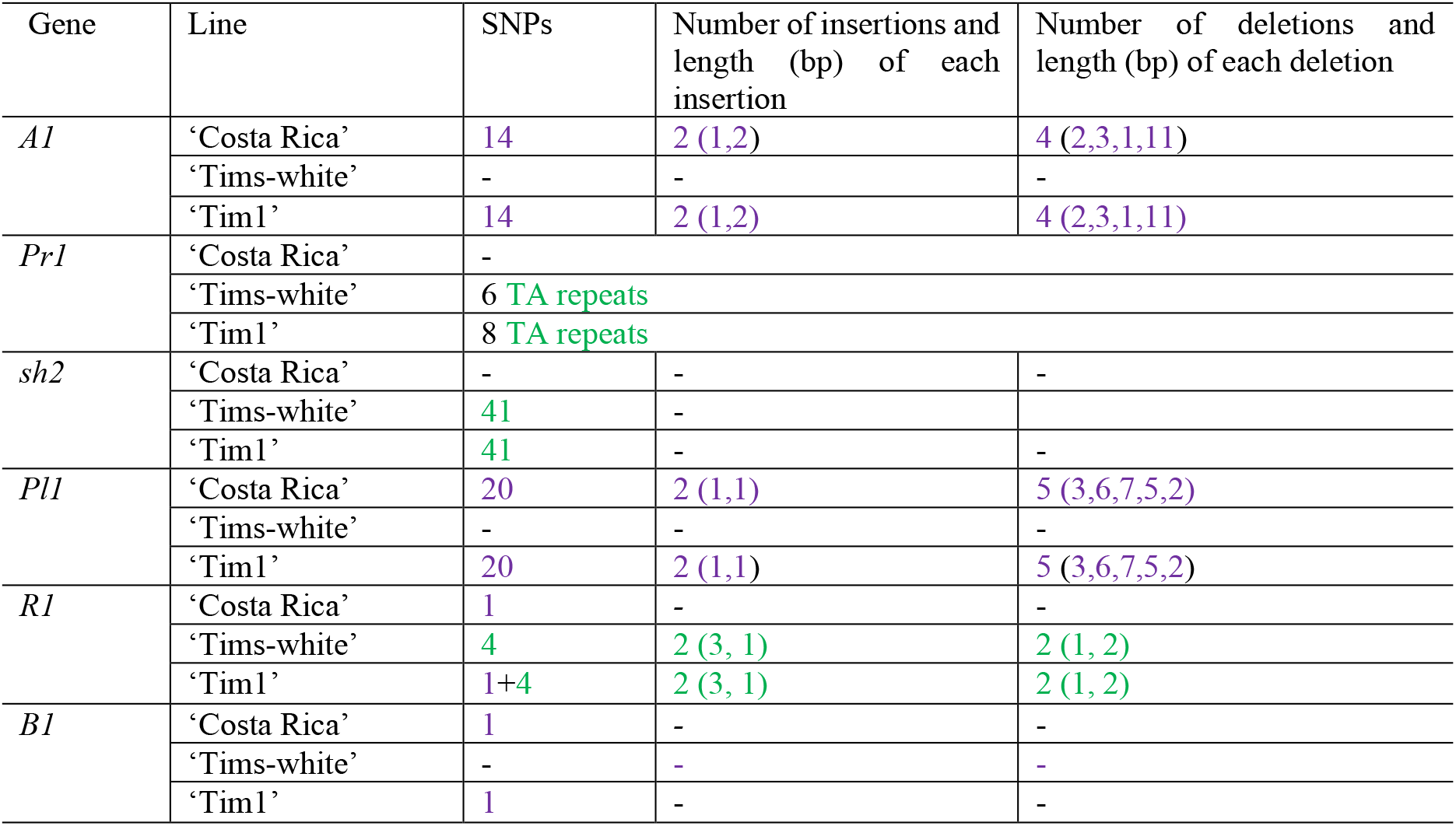
Summary of SNPs and InDels in the promoter region of target genes of ‘Tim1’ in common with ‘Costa Rica’ or/and ‘Tims-white’ parent. Purple colour indicates a common SNP/InDel in both ‘Tim1’ and ‘Costa Rica’. Green colour indicates a common SNP/InDel/repeat in both ‘Tim1’ and ‘Tims-white’.

There were 8-, and 6-dinucleotide (TA) repeats present in the promoter regions of the ‘Tim1’ and ‘Tims-white’ lines (Fig 3), whereas in the reference allele, there were 33-dinucleotide (TA) repeats present. The ‘Costa Rica’ line had no dinucleotide repeats.

**Fig 3.**
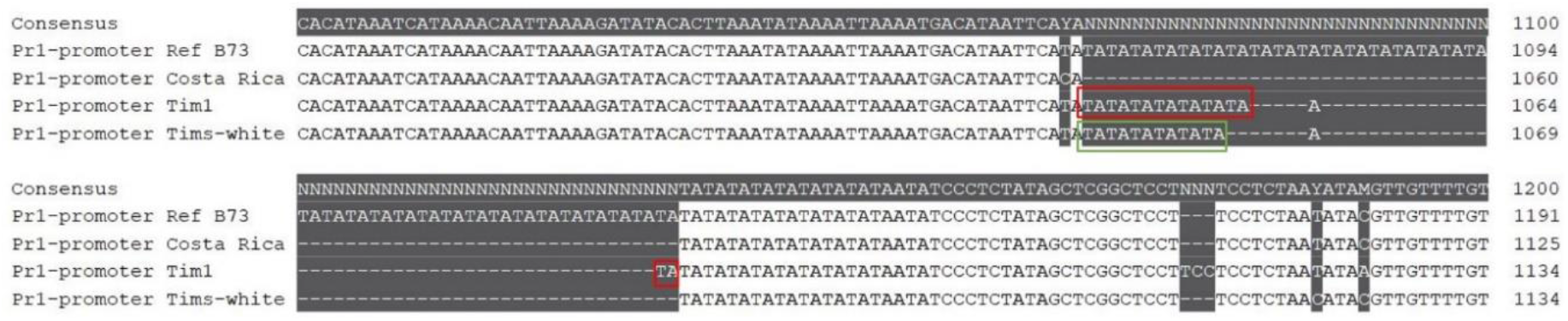
Dinucleotide (TA) repeats in the *pr1* gene present in ‘Tim1’ and ‘Tims-white’ lines. Red boxes indicate TA repeats in ‘Tim1’, and green box indicates TA repeats in ‘Tims-white’.

In case of *sh2* gene, 41 SNPs were present in the promoter region of both ‘Tims-white’ and ‘Tim1’ lines when they were compared with the wild type *Sh2* gene of the reference genome.

The *Pl1* gene included 20 SNPs, two insertions (each with 1 bp) and five deletions (3 bp, 6 bp, 7 bp, 5 bp and 2 bp) in the promoter region of both the ‘Costa Rica’ and ‘Tim1’ lines (Table 2) when compared with the reference genome, whereas the ‘Tims-white’ line was identical to the reference genome, again.

One heterozygous SNP was found in the CDS region of the *R1* gene both in ‘Costa Rica’ and ‘Tim1’ (Fig 2D), however in the promoter region there was one common SNP in both ‘Costa Rica’ and ‘Tim1’, and four common SNPs between ‘Tims-white’ and ‘Tim1’ (Table 2). Moreover, two insertions (3 bp and 1 bp) and two deletions (1 bp and 2 bp) were also present in both ‘Tims-white’ and ‘Tim1’.

There were one common SNP present both in the promoter region of *B1* gene of ‘Costa Rica’ and ‘Tim1’ (Table 2), while no SNP is present in ‘Tims-white’ line.

## Discussion

### Variation in the CDS regions

By breaking the close genetic linkage between the anthocyanin biosynthesis inactive allele, *anthocyaninless1 (a1)*, and super-sweet allele shrunken2 (*sh2)* novel purple-pericarp super-sweetcorn fixed line, ‘Tim1’ was obtained. The purple-pericarp maize ‘Costa Rica’ and white super-sweetcorn ‘Tims-white’ are the parental lines of ‘Tim1. Homozygous dominant anthocyanin biosynthesis allele, *A1* (*anthocyanin1*) and homozygous recessive sweetness allele, *sh2* were identified by only visual phenotyping based on purple colour development in the pericarp and shrunken appearance of the kernels at mature stage. In the current study, genomic study revealed the variation of anthocyanin biosynthesis structural and regulatory genes and sweetness structural gene in the CDS and promoter regions of the parent and novel progeny.

Previous studies by Muraya et al., (2015) and Yang et al., (2019) performed whole genome sequencing analysis based on homozygous/inbred lines. In this study, four ‘Costa Rica’ maize lines were used, and the sequenced ‘Costa Rica’ was a DNA mixture of these four lines. This was done to identify all potential variation that might exist in the ‘Costa Rica’ lines; due to the potential heterozygosity of certain genes associated with anthocyanin regulation.

A homozygous nonsynonymous SNP (Fig 1A) leading to the change of reference amino acid lysine to threonine (Lys12Thr) was observed both in purple maize, ‘Costa Rica’ parent and the developed purple-pericarp super-sweetcorn line, ‘Tim1’ in the anthocyanin biosynthesis structural gene, *A1*, and no nonsynonymous SNP was observed in white sweetcorn ‘Tims-white’ parent. This finding revealed that the ‘Costa Rica’ line is fixed (homozygous) for the functional anthocyanin biosynthesis dominant structural gene *A1*, enabling anthocyanin biosynthesis both in ‘Costa Rica’ and ‘Tim1’, although further study would be needed to confirm functionality. Previously, there has been no information regarding nonsynonymous SNPs in the *A1* gene, although study by Schwarz-Sommer et al., (1987) did indicate that frame-shift mutations invalidated functionality of the *A1* gene, and that one additional amino acid in the CDS region can still permit gene expression.

There was no common nonsynonymous SNP between ‘Tim1’ and ‘Costa Rica’ or ‘Tim1’ and ‘Tims-white’ in the case of the anthocyanin biosynthesis another structural gene *Pr1*. It may indicate that the promoter region is responsible for the change of colour from purple to red than the CDS region. A previous study by Sharma et al., (2011) with four *Pr1* alleles also found no difference between dominant and recessive genes in the CDS region.

One common homozygous non-synonymous SNP (Fig 1B) leading to the change of reference amino acid alanine to threonine (Ala52Thr) was observed both in the ‘Tims-white’ and ‘Tim1’ super-sweetcorn lines regarding sweetcorn structural gene *sh2*. However, no nonsynonymous SNP was observed in ‘Costa Rica’ parent. This finding is in agreement with recent research by Ruanjaichon et al., (2021), where a non-synonymous SNP change from C to T was considered the causal SNP for sweetness in corn. This result also confirms that ‘Tims-white’ and ‘Tim1’ are homozygous for the recessive sweetness allele, *sh2*, and the ‘Costa Rica’ line is homozygous for the dominant starch biosynthesis structural allele (*Sh2*).

Regarding the anthocyanin biosynthesis regulatory gene, *Pl1*, one homozygous (Fig 2A) and three heterozygous (Fig 2B) nonsynonymous SNPs were observed both in ‘Costa Rica’ and ‘Tim1’. Previous study by Cone et al., (1993) indicated that protein coding regions of dominant and recessive *pl* alleles were identical, which is in contrast with the current study. In addition, a six base pair deletion (Fig 2C) was also present in both ‘Costa Rica’ and ‘Tim1’. Earlier study by Yonemaru et al., (2018) found both SNPs and Indels in the exonic and intronic regions between a Peruvian purple maize cultivar and a yellow maize, however the results were not consistent with the current study.

Regarding *R1* gene, there was a common heterozygous non-synonymous SNP in ‘Costa Rica’ and ‘Tim1’ (Fig 2D). A heterozygous allele/ SNP may also impact on phenotype and be functional. This assumption is supported by the result of ‘Tims-white’ line, in which no SNP was observed compared to ‘Costa Rica’ and ‘Tim1’ in the same position for both the regulatory *Pl1* and *R1* genes. Therefore, *R1*, rather than *B1*, is the main bHLH transcription factor regulating pericarp anthocyanin along with the Myb transcription factor, *Pl1*.

Two common homozygous nonsynonymous SNPs in the ‘Tims-white’ and ‘Tim1’ lines, and no nonsynonymous SNP in ‘Costa Rica’ were observed in case of *B1* gene (Fig 2E). As there was no nonsynonymous SNP in the purple maize line ‘Costa Rica’, it may be inferred that *B1* may not have role in pericarp anthocyanin regulation, in contrast to the earlier reports by Lago et al., 2014; and Lago et al., 2012. Recent studies by Chatham and Juvik, (2021), Chatham et al., (2019), and Portwood et al., (2019) have been reported that *R1* instead of *B1*, play a regulatory role as a bHLH transcription factor in pericarp anthocyanin biosynthesis.

### Variation in the promoter regions

There were 14 SNPs, two insertions and four deletions common to ‘Costa Rica’ and ‘Tim1’ in the promoter region of the *A1* gene (Table 2). A previous study by Tuerck and Fromm, (1994) also indicated that the promoter region of the *A1* gene of maize is critical for this anthocyanin biosynthesis gene. However, there was no SNP or InDel based analysis presented in that study. Moreover, the alleles of the *A1* gene were different to those found in the present study. Another study by Bernhardt and Stich, (1988), however, found a second functional *A1* gene in maize.

8-, 6-, and 33-dinucleotide (TA) repeats were present in the promoter regions of the *Pr1* gene of ‘Tim1’, ‘Tims-white’, and reference allele, respectively (Fig 3). Previous research by Sharma et al., (2011) found a 24-dinucleotide (TA) repeat in a mutant *pr1* alleles and were considered the reason for the defective nature of recessive *pr1* allele. There were no dinucleotide repeats present in ‘Costa Rica’ parent, which implies that a functional *Pr1* is active the ‘Costa Rica’ line.

41 SNPs both in the promoter region of ‘Tims-white’ and ‘Tim1’ lines regarding the *sh2* gene indicate their identical nature in the promoter region, as like CDS region.

In case of *Pl1* gene, 20 SNPs, two insertions and five deletions were observed common in between ‘Costa Rica’ and ‘Tim1’ lines (Table 2) in the promoter region. Previous study by Cone et al., (1993) indicated that although the protein coding regions of the dominant and recessive *Pl* alleles were similar, they had sequence differences in the promoter region. Another study by Pilu et al., (2003) revealed that the *pl-bol3* gene, which also regulates anthocyanin biosynthesis, had sequence differences in the promoter region compared to the dominant allele, *Pl-Rh*.

Regarding *R1* gene, there was one common SNP in both ‘Costa Rica’ and ‘Tim1’, and four SNPs, two insertions and two deletions were present in both ‘Tims-white’ and ‘Tim1’ (Table 2). In a previous study by Consonni et al., (1993) similarity as well as differences among different *R* genes (S, Sn, B-Peru and Lc) was also reported.

In conclusion, the novel purple-pericarp super-sweetcorn line, ‘Tim1’, and its purple-pericarp parent, ‘Costa Rica’, have functional nonsynonymous SNPs in common within the CDS region of the *A1, Pl1* and *R1* genes, required for anthocyanin biosynthesis. In contrast, ‘Tim1’ had nonsynonymous SNPs in the CDS region of *B1* in common with its white super-sweetcorn parent, ‘Tim’s-white’, indicating that *B1* may not play a role in pericarp anthocyanin regulation. Regarding the sweetcorn structural gene *sh2*, ‘Tim1’ had non-synonymous SNPs in the CDS region in common with that observed in ‘Tim’s-white’, the super-sweetcorn parent. This SNP is functional as a sweetcorn gene and stops formation of starch. In the promoter region, the *sh2* gene showed similarity between ‘Tim1’ and ‘Tims-white’, again. In the promoter region, similarity in the case of SNPs, InDels between ‘Tim1’ and ‘Costa Rica’ regarding the *A1, Pl1, R1* and *B1* genes was present. However, in the case of the *R1* gene, partial similarity of SNPs and InDels between ‘Tim1’ and ‘Tims-white’ lines were also observed. Both in the ‘Tim1’ and ‘Tims-white’ lines, the *Pr1* gene had dinucleotide (TA) repeats in the promoter region, which may be the cause of the defective (redder colour in ‘Tim1’ than purple in ‘Costa Rica’) nature of the recessive *pr1* allele.

## Materials and methods

### Plant materials

Purple-pericarp starchy (non-sweet) maize (‘Costa Rica’, *A1Sh2*.*A1Sh2*) parental line, white super-sweetcorn parental line (‘Tims-white’, *a1sh2*.*a1sh2*) and developed F6 purple-pericarp super-sweetcorn line, ‘Tim1’ (*A1A1*.*sh2sh2*) were used in this experimental work (Fig 4A-F). Flow chart related to the development of F6 purple-pericarp super-sweetcorn line, ‘Tim1’ (*A1A1*.*sh2sh2*) illustrates all field experiments (Fig 4G).

**Fig 4.**
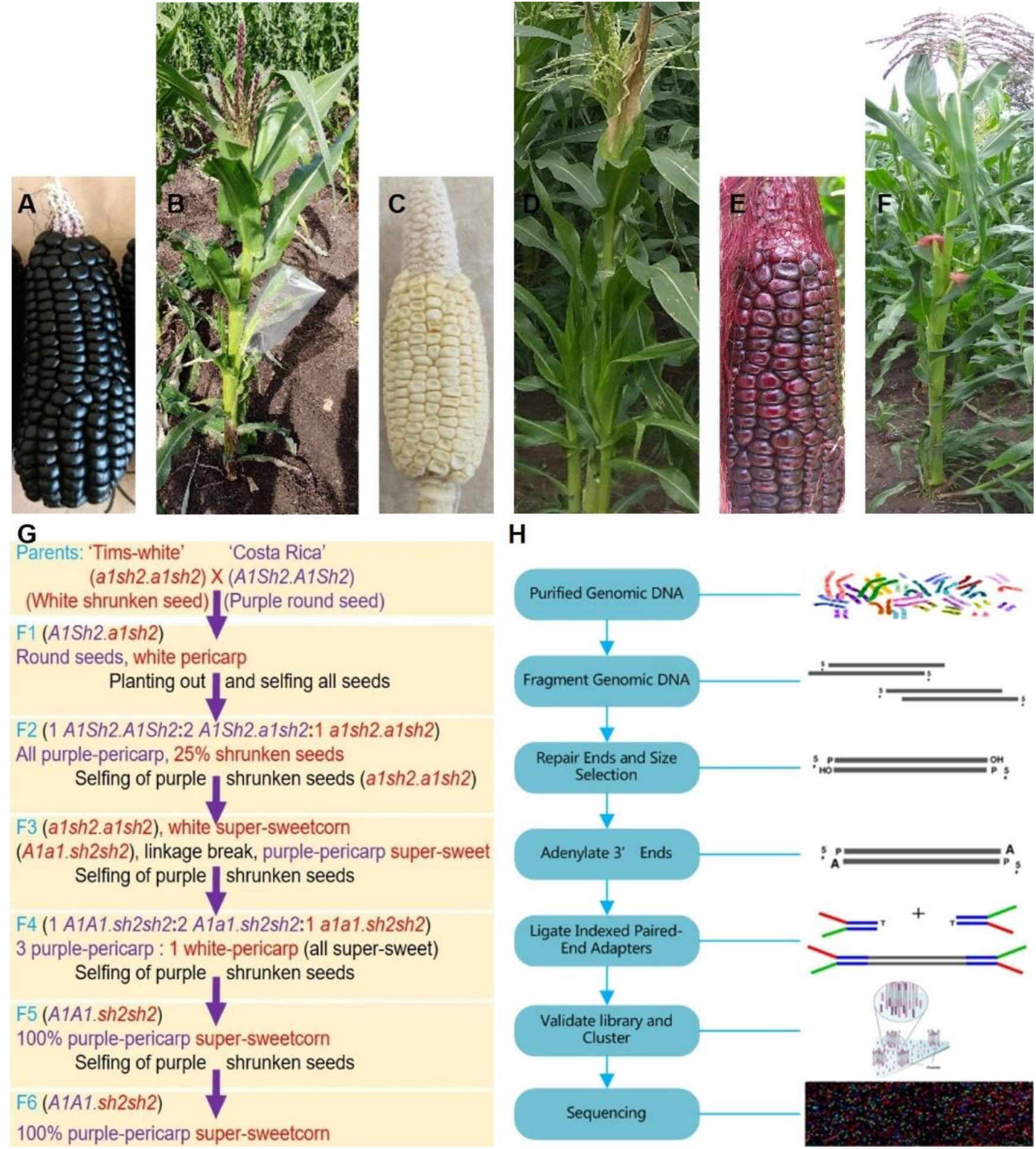
Seeds and plants of ‘Costa Rica’, ‘Tims-white’, and ‘Tim1’ used in field experiments as well as DNA re-sequencing: **A)** ‘Costa Rica’ cob with purple-pericarp starchy (non-sweet) round kernels, **B)** ‘Costa Rica’ plant, **C)** ‘Tims-white’ cob with white-pericarp non-starchy (sweet) kernels, **D)** ‘Tims-white’ plant, **E)** ‘Tim1’ cob with purple-pericarp non-starchy (sweet) kernels, **F)** ‘Tim1’ plant, **G)** Flow chart showing the purple-pericarp super-sweetcorn development scheme from the initial cross to the development of the F6 progeny; **H)** DNA re-sequencing by Illumina NovaSeq.

### DNA extraction of four ‘Costa Rica’ lines and ‘Tims-white’ and ‘Tim1’ lines for re-sequencing

DNA of four ‘Costa Rica’ lines was extracted individually, equal concentration (35 ng/ul) of DNA from each line was pooled together. This pooled DNA was sequenced to attain variation information within the coding and promoter regions of the target genes. As ‘Tims-white’ line was an inbred line, and ‘Tim1’ was a developed F6 generation self-pollinated line, no pooling of DNA was necessary.

Extraction of DNA was performed with 14-day-old leaf samples of the four purple-pericarp maize lines (‘Costa Rica’), one white super-sweetcorn line (‘Tims-white’) and one developed F6 purple-pericarp super-sweetcorn line (‘Tim1’) as a earlier method by Vivekananda et al., (2018), with some modifications, as follows.

At first, 100 mL extraction buffer (50 mM Tris-HCl, 25 mM EDTA, 1% SDS, 300 mM NaCl) stock was prepared with 5 mL of 1 M Tris-HCl pH 8.0, 5 mL of 0.5 M EDTA pH 8.0, 21.43 mL of 1.4 M NaCl, 10 mL of 10% SDS, and added required volume (58.57 mL) of DH20.

Before DNA extraction, leaf samples (in falcon tubes) were removed from -80 ⁰C and the tubes were put into liquid N2 container. 200 μL of extraction buffer (50 mM Tris-HCl, 25 mM EDTA, 1% SDS, 300 mM NaCl) was put into 2 mL microcentrifuge tube. Some 1 cm2 sizes leaf tissues were transferred to a 2 mL microcentrifuge tube and ground using an Eppendorf micro pestle (Sigma Aldrich). An additional 200 μL of extraction buffer was added to the tube and vortexed for 10 sec. The tubes were subsequently incubated in a water bath at 65 ⁰C for 30 min. The tube contents were then mixed well by inversion and cooled to ambient room temperature (25 °C). 200 μL of chloroform:isoamyl alcohol (24:1) and 200 μL of Tris-saturated phenol was added and mixed by inverting the tube and vortexing for 10 sec. The mixture was centrifuged at 13,000 rpm for 15 min. 600 uL supernatant (upper aqueous layer) was then transferred into a new 2 mL tube.

2 uL of 50 mg/mL (5 ul of 20 mg/mL) RNaseA was added to the tube, mixed well by inversion, and incubated at 65 ⁰C for 30 min. 600 μL of chloroform was added and mixed gently by inversion for up to 1 min, and the samples centrifuged at 13,000 rpm for 15 min. The upper aqueous layer (600 uL) was again transferred to a new 2 mL tube. 600 μl of ice-cold isopropanol was added and mixed gently by inverting the tube (1-30 min) until strings of DNA were observed. If strings of DNA were not observed, it was transferred to a -20 ⁰C freezer for 15 min and rechecked for string formation.

The mixture was then centrifuged at 13,000 rpm for 15 min. The supernatant was discarded. The pellet was washed with 800 uL ice-cold 70% ethanol and mixed gently by inversion for 1 min. The tube was again centrifuged at 13,000 rpm for 15 min. The ethanol solution was poured off and the tube placed on a drying rack and air-dried for about 5 min. 50 μL of TE buffer or sterile DH2O was added to re-suspend the DNA pellet, which was mixed gently to allow the release of the bound DNA into the buffer.

DNA quality was measured using the spectrometry absorbance ratios of 260/280 nm and 260/230 nm (NanoDrop, Thermofisher Scientific) respectively, which are used for primary and secondary contamination assessment. The extracted DNA was then run on agarose gel electrophoresis. The quantity of DNA was measured using lambda DNA standards. The 260/280 absorbance ratio between 1.8-2.0, and the 260/230 absorbance ratio between 2.0-2.2, and non-shearing of DNA were regarded as good quality DNA. Then, re-sequencing of DNA (Fig 4H) was performed by Illumina NovaSeq (Genewiz, China).

### Variant analysis at gene level using CLC Genomics Workbench

Homozygous and heterozygous SNPs, synonymous and nonsynonymous SNPs in the CDS region of each of the target gene were analysed against the annotated reference yellow starchy maize B73 (NAM5.0) genome (Woodhouse et al., 2021) by applying CLC Genomics Workbench 21.0.4 of Qiagen (http://www.clcbio.com).

## Availability of data and materials

Sequence data from this article have been deposited with the NCBI (https://www.ncbi.nlm.nih.gov/) Data Libraries under BioProject accession number **PRJNA860024**.

## Author contributions

AA performed the field experiments, DNA extraction, and variant analysis. AA composed the full manuscript; TJO supervised the PhD project, and AKM and RJH co-supervised. AKM, RJH and TJO reviewed and edited the manuscript. AA updated the manuscript. All authors read and approved the final manuscript.

## Acknowledgements

This article is the part of PhD research of Apurba Anirban. AA is grateful to the University of Queensland for providing with PhD Fellowship.

